# Rise in mortality involving poisoning by medicaments other than narcotics, including poisoning by psychotropic drugs in different age/racial groups in the US

**DOI:** 10.1101/509729

**Authors:** Edward Goldstein

## Abstract

**Background:** Increase in mortality involving poisoning, particularly by narcotics, is known to have been one of the factors that affected life expectancy in the US during the last two decades, especially for white Americans and Native Americans. However, the contribution of medicaments other than narcotics to mortality in different racial/age groups is less studied.

**Methods:** We regressed annual rates of mortality involving poisoning by medicaments but not narcotics/psychodysleptics (ICD-10 codes T36-39.xx or T41-50.8 but not T40.xx present as either underlying or contributing causes of death), as well as annual rates of mortality for certain subcategories of the above, including mortality involving poisoning by psychotropic drugs but not narcotics/psychodysleptics (ICD-10 codes T43.xx but not T40.xx present as either underlying or contributing causes of death) in different age/racial groups for both the 2000-2011 period and the 2011-2017 period against calendar year.

**Results:** Annual numbers of deaths involving poisoning by medicaments but not narcotics/psychodysleptics grew from 4,332 between 2000-2001 to 11,401 between 2016-2017, with the growth in the rates of those deaths being higher for the 2011-2017 period compared to the 2000-2011 period. The largest increases in the rates of mortality involving poisoning by medicaments but not narcotics/psychodysleptics were in non-elderly Non-Hispanic Native Americans, followed by Non-Hispanic whites. Most of those increases came from increases in the rates of mortality involving poisoning by psychotropic medications; the latter rates grew for the period of 2015-2017 vs. 2000-2002 by factors ranging from 2.75 for ages 35-44y to 5.37 for ages 55-64y.

**Conclusions:** There were major increases in mortality involving poisoning by non-narcotic, particularly psychotropic medicaments, especially in non-elderly non-Hispanic whites and Native Americans. Our results, and the epidemiological data on mortality involving poisoning by different drugs and medications in the US, which are quite different from the ones in many other countries support the need for a comprehensive evaluation of the effect of various drugs, including psychotropic medications on health-related outcomes, the associated mortality the does not involve poisoning being listed on a death certificate, and the impact of medication misuse.

## Introduction

During the last two decades, life expectancy in white Americans has been decreasing compared to other major racial groups. By 2006, life expectancy in Hispanics became about 2.1 years higher than in non-Hispanic whites; by 2014, that gap has grown to about 3.2 years (Figure 4 in ref. [1]). Life expectancy in African Americans is consistently lower than in whites, though the gap has been decreasing significantly in the recent years (Figure 4 in ref. [2]; Figure 4 in ref. [1]). In absolute terms, life expectancy in whites has been declining starting 2014 [1,3], together with the overall decline in life expectancy in the US starting 2015 [4] amid an ongoing epidemic of drug overdose deaths [5,6].

Several factors behind the relative decline in life expectancy in white Americans have been proposed, [4,7]. In particular, as suggested by a study that compared midlife mortality for the 2013-2015 period vs. the 1999-2001 period [8], increases in mortality by poisoning, suicide, and liver disease in non-elderly white Americans were notably higher compared to Hispanics and blacks. Moreover, those increases, particularly for deaths by poisoning, took place in both the urban/suburban and rural areas (Figure 1 in ref. [8]). While the use of narcotics [5,6] is known to be a contributor to the long-term trends in the rates of midlife mortality by poisoning and drug overdose [9,8], misuse of medicaments other than narcotics may also have a pernicious health-related effect, including contribution to mortality in different population groups, particularly white Americans and Native Americans [9]. We note that the rates of consumption of certain medicaments in white Americans are higher than in other racial groups (excluding Native Americans) [10]. In particular, there are major differences in the rates of consumption of antibiotics [11,12], psychotropic drugs [13], including antidepressants [14,15], and sedative-hypnotic medications [16–18] between white Americans and other racial groups in the US except Native Americans. A number of studies, including meta-analyses, suggest elevated risks for mortality, including sudden cardiac death (SCD) associated with the use of various psychotropic drugs (both antipsychotics and antidepressants), as well as certain sedative-hypnotic drugs (benzodiazepines) [19–24], though firmly establishing causal links in such studies can be challenging [25]. For the case of antibiotics, recent work suggests an association between prescribing rates for certain antibiotics, particularly penicillins in the elderly, and mortality and hospitalization rates for sepsis/septicemia [26,27], with that association presumably being mediated by antimicrobial resistance [28]. Further work is needed to gain a more comprehensive understanding of the effect of prescribing and misuse of different medications on health-related, including mortality outcomes in different age/racial groups.

In this paper, we study one of the more direct types of contribution of medication use to mortality, namely mortality that involves poisoning by medications. We note that the rise in the rates of mortality associated with narcotic, including opiate overdose is well-documented, e.g. [5,6,9]. However, trends in mortality involving poisoning by various medicaments other than narcotics in the US are apparently not well studied. Here, we utilized the US CDC Wonder mortality database [29] to examine trends in mortality involving poisoning by medicaments other than narcotics/psychodysleptics in different age/racial groups in the US, including trends in mortality rates for certain subcategories of those deaths, particularly deaths involving poisoning by psychotropic, as well as sedative-hypnotic drugs. Our aim is to characterize the volume of such deaths, as well as trends in the rates of those deaths in different age/racial groups during different time periods. In the Discussion section, we’ll mention the comparison with the corresponding trends in other countries and the impact of medication misuse. This initial analysis is meant to provide rationale for further study of the contribution of medication use to mortality outcomes, including mortality that does not involve poisoning by medicaments present on the death certificate (e.g. [19–24;26,27]).

## Methods

### Data

We have extracted annual mortality data between 2000-2017 stratified by age group (25-34y, 35-44y, 45-54y, 55-64y, 65-74y) and racial group (Hispanics, Non-Hispanic whites, Non-Hispanic blacks, Non-Hispanic Asian-Americans, Non-Hispanic Native Americans) from the US CDC Wonder database [29]. While the 1999 data are also available in ref. [29], the 1999 mortality rates in the various categories that we have considered are generally higher than those rates during the subsequent years, possibly having to do with the transition from ICD-9 to ICD-10 coding around that time [30] and other factors relating to coding; correspondingly, the 1999 data were excluded from our analyses. The mortality data between 2000-2017 were extracted for five types of deaths:

Type 1. Deaths involving poisoning by drugs, medicaments, and other biological substances: ICD-10 codes T36-50.xx present as either underlying or contributing causes of death
Type 2. Deaths involving poisoning by narcotics and psychodysleptics: ICD-10 codes T40.xx present as either underlying or contributing causes of death
Type 3. Deaths involving poisoning by specific drugs, medicaments, and other biological substances: ICD-10 codes T36-50.8 present as either underlying or contributing causes of death. We note that ICD-10 codes T50.9 represent poisoning by other and unspecified drugs, medicaments, and other biological substances, and those deaths are not included in type 3.
Type 4. Deaths involving poisoning by either psychotropic drugs (antipsychotic/neuroleptic drugs and antidepressants) or narcotics and psychodysleptics: ICD-10 codes T43.xx or T40.xx present as either underlying or contributing causes of death
Type 5. Deaths involving poisoning by either sedative-hypnotic drugs (barbiturates and benzodiazepines) or narcotics and psychodysleptics: ICD-10 codes T42.3-43.4 or T40.xx present as either underlying or contributing causes of death

Those data were used to calculate the mortality rates for different categories of death used in this study. Those categories, and the calculation of the corresponding mortality rates are described in Table 1 below.

**Table 1:** Different categories of death used in the analyses, and the relevant ICD-10 reflecting either the underlying or contributing causes of death.

| Category | Description | ICD-10<br>(underlying or<br>contributing) | Extraction process |
| --- | --- | --- | --- |
| A | Poisoning by drugs,<br>medicaments, and other<br>biological substances | T36-50.xx | Rates in type 1 in the<br>Methods |
| B | Poisoning by narcotics and<br>psychodysleptics | T40.xx | Rates in type 2 in the<br>Methods |
| C | Poisoning by medicaments but not narcotics and psychodysleptics | T36-39.xx or T41-50.8 but not T40.xx | Rates in type 2 subtracted from rates in type 3 in the Methods |
| D | Poisoning by psychotropic drugs but not narcotics and psychodysleptics | T43.xx but not T40.xx | Rates in type 2 subtracted from rates in type 4 in the Methods |
| E | Poisoning by sedative-hypnotic drugs but not narcotics and psychodysleptics | T42.3-42.4 but not T40.xx | Rates in type 2 subtracted from rates in type 5 in the Methods |

We note that the reason behind the subtraction for categories C-E is that if one wants to find the number of deaths that have causes X but not Y (e.g. psychotropic medications but not narcotics/psychodysleptics) listed on the death certificate, one subtracts the number of deaths that have causes Y listed on a death certificate from the number of deaths that have either causes X or causes Y listed on a death certificate.

### Trends in mortality rates (regression analysis)

For each age/racial group, we have regressed the corresponding annual rates of mortality for categoriess A through E (Table 1) against the calendar year covariate (univariate regression). Given the apparent change in the trends for mortality involving poisoning by medicaments (including psychotropic drugs) but not narcotics and psychodysleptics in certain age/racial groups starting around 2012 (Figures 3, 4), the regression analyses were performed separately for the 2000-2011 period and the 2011-2017 period. The regression coefficient for the calendar year covariate estimates the annual increase in the rate of mortality for the corresponding category (A. through E. in Table 1) in the given age/racial group during the corresponding time period (2000-2011 or 2011-2017).

### Relative changes in mortality rates

In addition to the evaluation of trends in mortality for the different categories of death (listed in Table 1) in different age/racial groups, we also calculated the relative (fold) increases in the rates of mortality for categories B, C, D, E in Table 1 in different age groups of US adults. We considered, for different age groups of adults and categories B through E, changes in average annual mortality rates for the 2015-2017 period vs. 2000-2002 period, as well as for the 2011-2014 period vs. 2000-2002 period (with the latter relative change evaluated to allow for the comparison with changes in prescribing rates for different medications during the same period recorded in ref. [31]).

### Spatial variability in the rates of mortality involving poisoning by different substances

In addition to temporal changes in the rates of mortality involving poisoning by different substances, we also looked at state-specific rates of mortality involving poisoning by psychotropic drugs but not narcotics/psychodysleptics (category D in Table 1), as well as mortality involving poisoning by narcotics/psychodysleptics (category B in Table 1), including correlations between those state-specific rates.

## Results

Figures 1–5 plot the annual rates of mortality (per 100,000 individuals) between 2000-2017 for the different categories of death (A through E) in Table 1. For the younger age groups (ages 25-34y, 35-44y and 45-54y), the largest increases in mortality rates for all the categories A-E were in non-Hispanic Native Americans, followed by non-Hispanic whites. Figure 1 plots the annual rates of mortality involving poisoning by drugs, medicaments, and other biological substances (category A in Table 1) between 2000-2017 for the different age/racial groups. Figure 2 plots the annual rates of mortality involving poisoning by narcotics and psychodysleptics (category B in Table 1) between 2000-2017 for the different age/racial groups. We note that most deaths involving poisoning by drugs, medicaments, and other biological substances are deaths involving poisoning by narcotics and psychodysleptics (Figure 2 vs. 1). Rates of mortality involving poisoning by narcotics and psychodysleptics in white and Native Americans aged under 55y are high. For example, for Non-Hispanic whites, by 2017 about a third of all deaths in individual aged 25-34y, and about 20% of all deaths in individuals aged 35-44y involved poisoning by narcotics and psychodysleptics. The largest rises in mortality rates involving poisoning by narcotics and psychodysleptics in older age groups (55-64y and 65-74y), particularly during the most recent years, were in non-Hispanic blacks (Figure 2).

**Figure 1:**
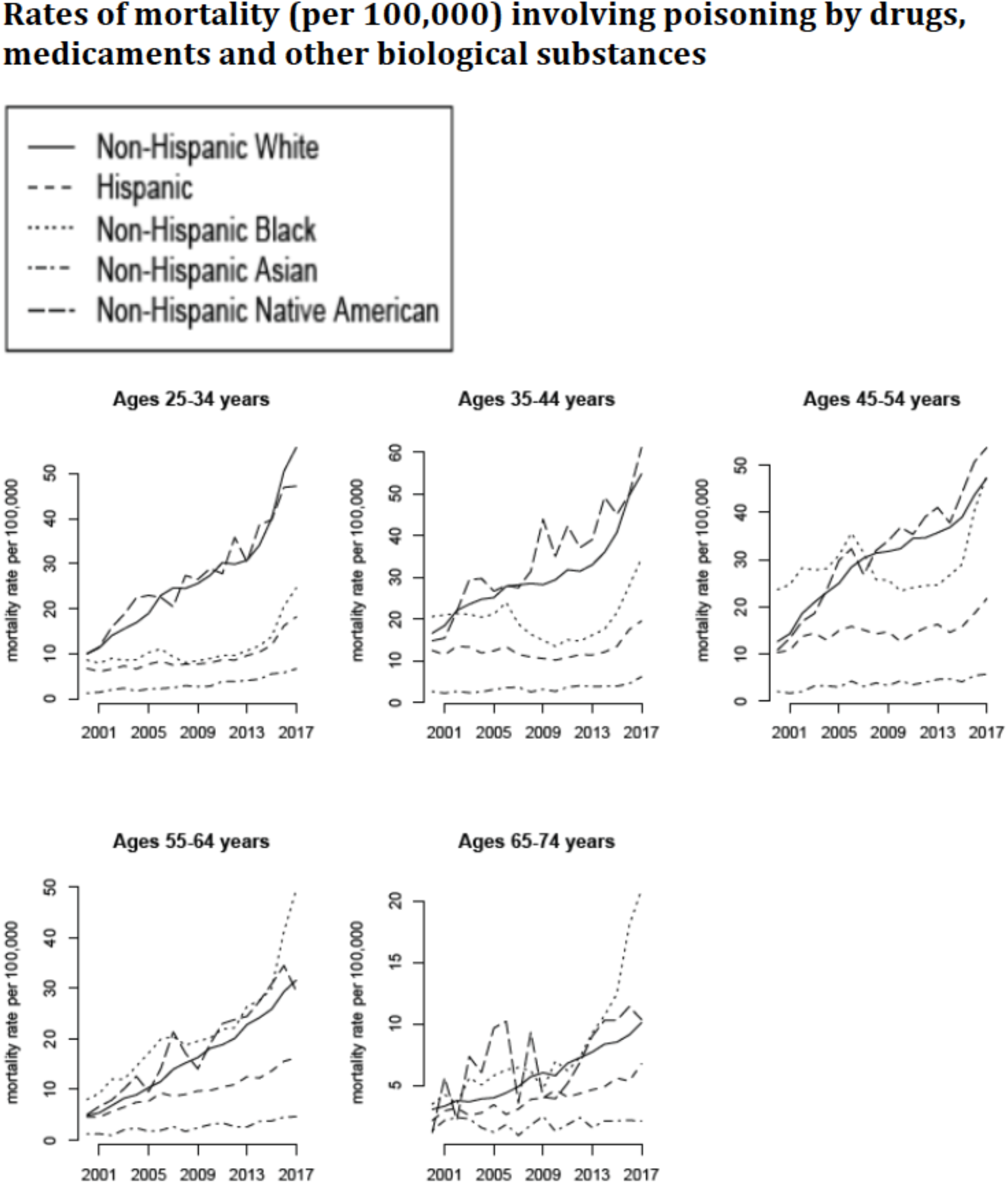
Annual rates of mortality involving poisoning by drugs, medicaments and other biological substances (ICD-10 codes T36-50.xx present as either underlying or contributing causes of death; category A in Table 1) between 2000-2017 per 100,000 individuals in different age/racial groups.

**Figure 2:**
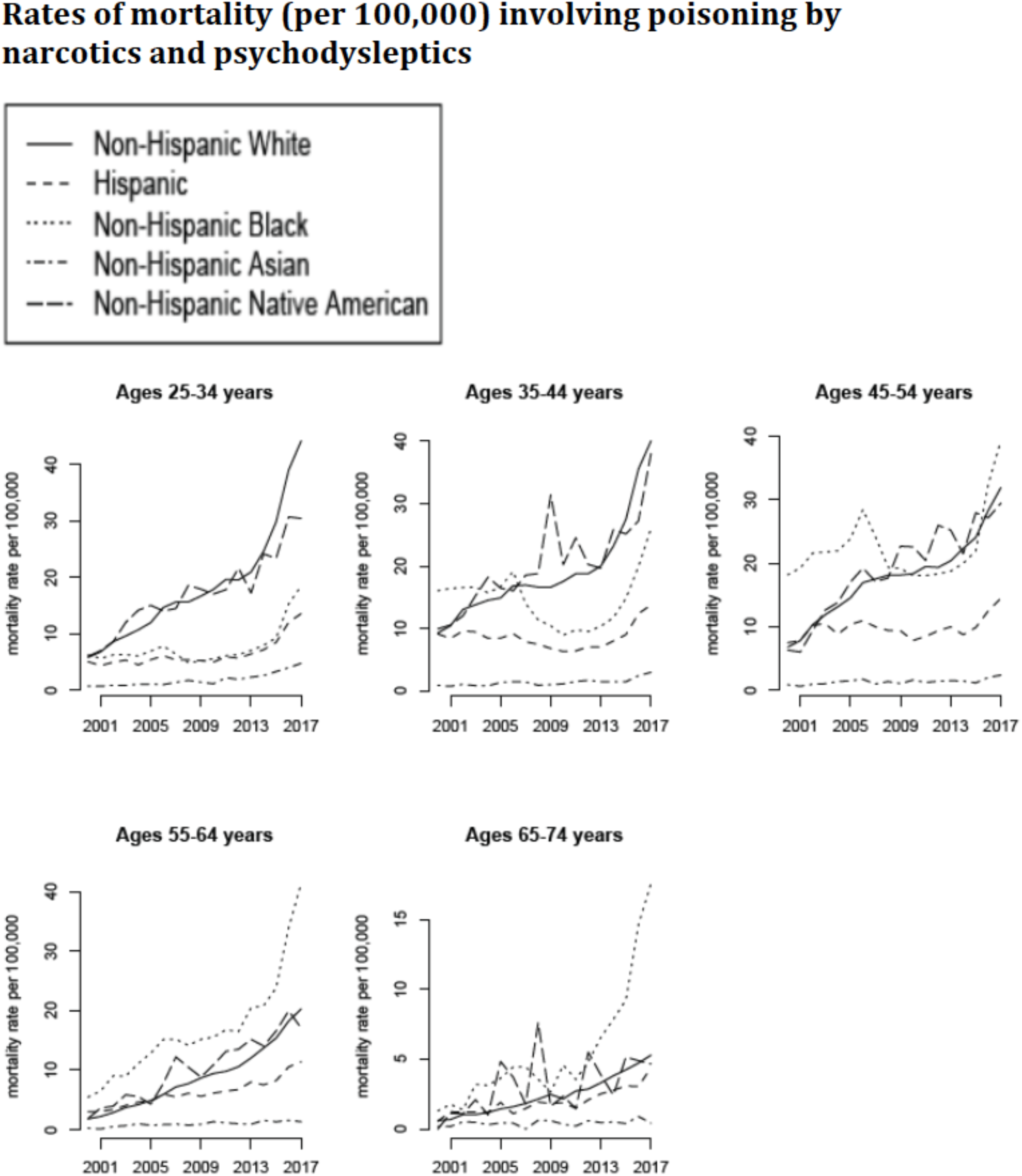
Annual rates of mortality involving poisoning by narcotics and psychodysleptics (ICD-10 codes T40.xx present as either underlying or contributing causes of death; category B in Table 1) between 2000-2017 per 100,000 individuals in different age/racial groups.

The number of deaths involving poisoning by medicaments but not narcotics and psychodysleptics in the US increased significantly during our study period, from the annual average of 4,332 between 2000-2001 to the annual average of 11,401 between 2016-2017. Figure 3 plots the annual rates of mortality involving poisoning by medicaments but not narcotics and psychodysleptics (category C in Table 1) between 2000-2017 for the different age/racial groups. We note that those rates are lower than the rates of mortality involving poisoning by narcotics and psychodysleptics (Figure 3 vs. Figure 2). Figures 4 and 5 plot the annual rates of mortality involving poisoning by psychotropic drugs but not narcotics and psychodysleptics (category D in Table 1) and sedative-hypnotic drugs but not narcotics and psychodysleptics (category E in Table 1) between 2000-2017 for the different age/racial groups. The majority of deaths involving poisoning by medicaments but not narcotics and psychodysleptics, as well as most of the increases in those death rates were for deaths involving poisoning by psychotropic drugs but not narcotics and psychodysleptics (Figure 3 vs. 4).

**Figure 3:**
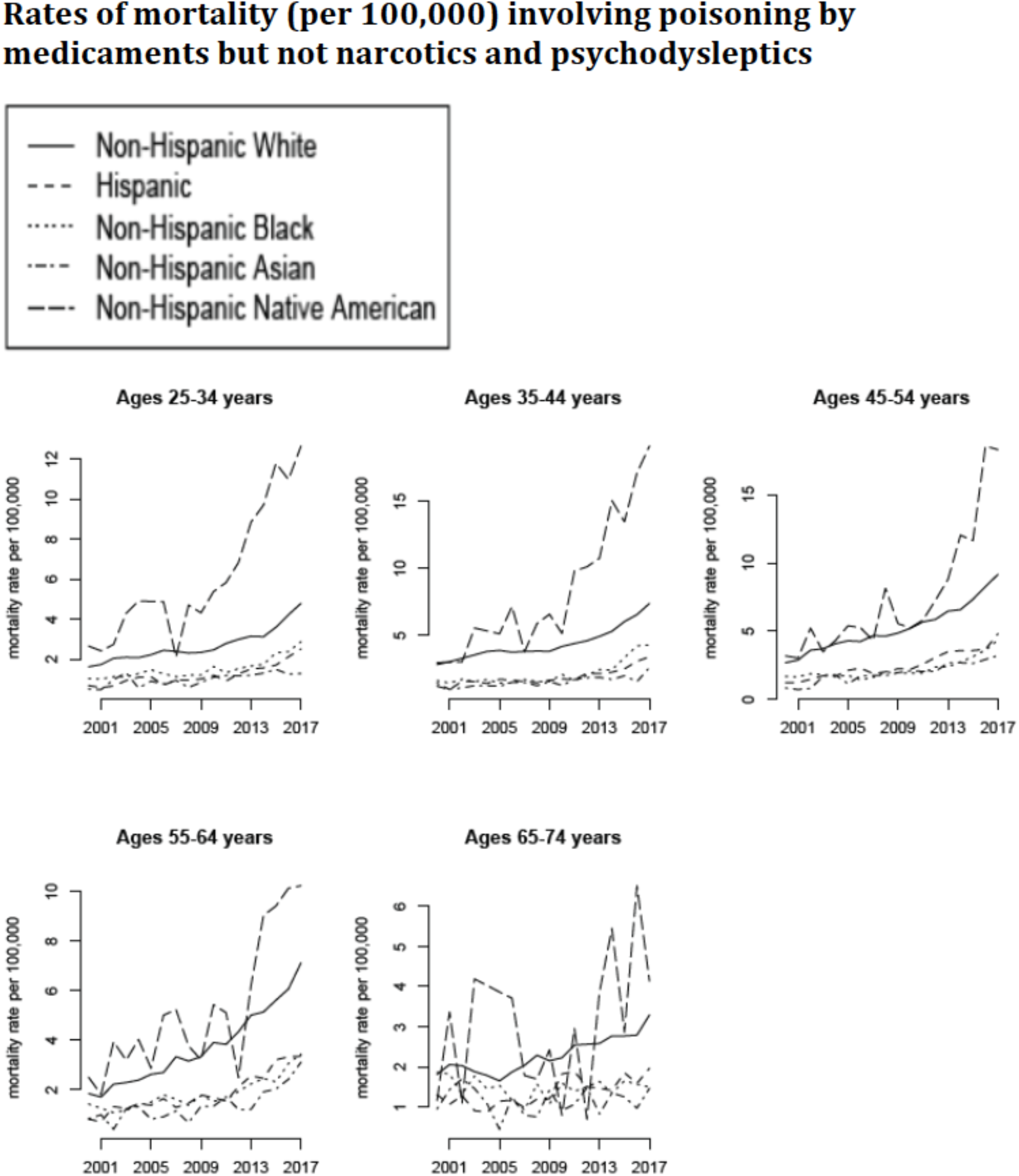
Annual rates of mortality involving poisoning by medicaments but not narcotics and psychodysleptics (ICD-10 codes T36-39.xx or T41-50.8 but not T40.xx not present as either underlying or contributing causes of death; category C in Table 1) between 2000-2017 per 100,000 individuals in different age/racial groups.

**Figure 4:**
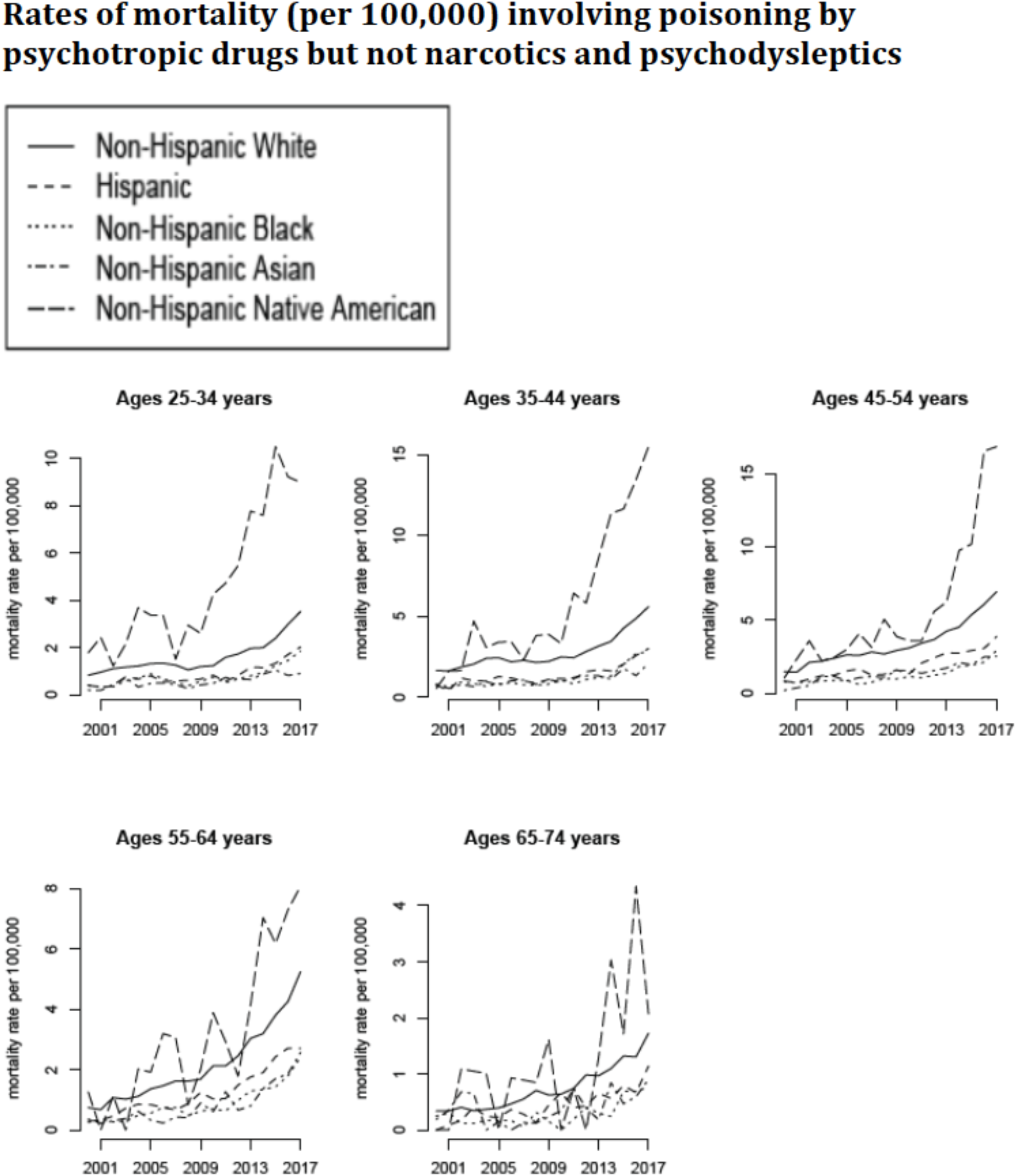
Annual rates of mortality involving poisoning by psychotropic drugs but not narcotics and psychodysleptics (ICD-10 codes T43.xx but not T40.xx not present as either underlying or contributing causes of death; category D in Table 1) between 2000-2017 per 100,000 individuals in different age/racial groups.

**Figure 5:**
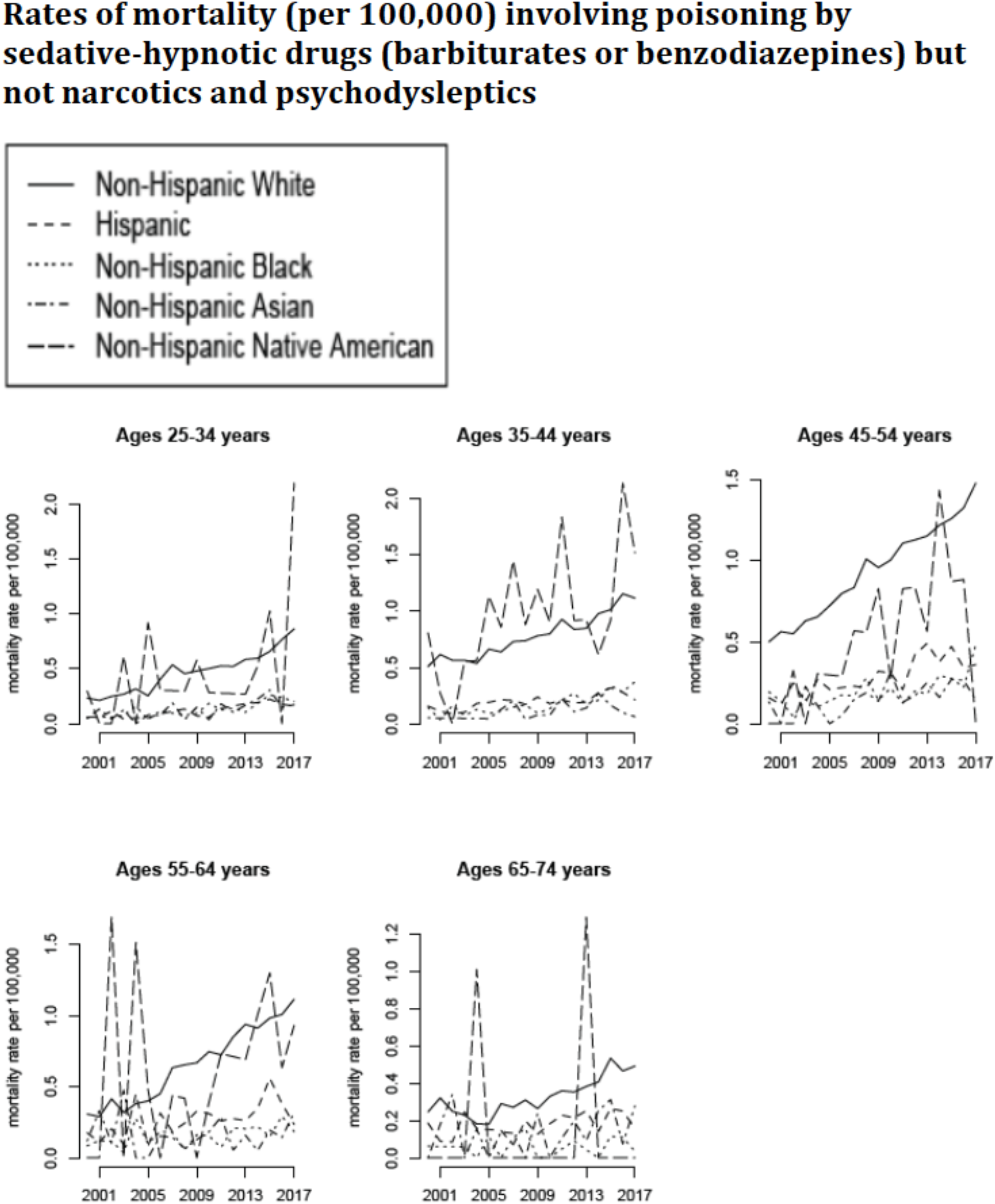
Annual rates of mortality involving poisoning by sedative-hypnotic drugs but not narcotics and psychodysleptics (ICD-10 codes T42.3-42.4 but not T40.xx not present as either underlying or contributing causes of death; category E in Table 1) between 2000-2017 per 100,000 individuals in different age/racial groups.

Figures 3 and 4 suggest an apparent change in the trends for mortality involving poisoning by medicaments (including psychotropic drugs) but not narcotics and psychodysleptics in certain age/racial groups (particularly Non-Hispanic Native Americans, Non-Hispanic whites and Hispanics aged 25-44y) starting around 2012. Table 2 gives the estimates of the annual increase in the rate of mortality involving poisoning by medicaments but not narcotics and psychodysleptics in different age/racial groups between 2000-2011, as well as between 2011-2017 (see the *Trends in mortality rates (regression analysis)* subsection of the Methods). Annual increases in the rates of mortality involving poisoning by medicaments but not narcotics and psychodysleptics between 2011-2017 are significantly higher than the corresponding increases during the 2000-2011 period, with the largest relative change between the two time periods being in Non-Hispanic blacks. The greatest increases in the rates of mortality involving poisoning by medicaments but not narcotics and psychodysleptics during each time period are in non-elderly Non-Hispanic Native Americans, followed by Non-Hispanic whites.

**Table 2:** Annual change in mortality rates involving poisoning by medicaments but not narcotics and psychodysleptics (ICD-10 codes T36-39.xx or T41-50.8 but not T40.xx not present as either underlying or contributing causes of death; category C in Table 1) between 2000-2011 and between 2011-2017 per 100,000 individuals in different age/racial groups

| **2000-2011 period** |  |  |  |  |  |
| --- | --- | --- | --- | --- | --- |
|  | Ages<br>25-34y | Ages<br>35-44y | Ages<br>45-54y | Ages<br>55-64y | Ages<br>65-74y |
| Non-Hispanic<br>whites | **0.08**<br>(0.06,0.1) | **0.11**<br>(0.08,0.14) | **0.24**<br>(0.21,0.27) | **0.20**<br>(0.17,0.23) | **0.05**<br>(0.02,0.08) |
| Hispanics | **0.03**<br>(0.01,0.06) | **0.05**<br>(0.02,0.07) | **0.11**<br>(0.08,0.14) | **0.07**<br>(0.04,0.11) | **0.05**<br>(0.01,0.1) |
| Non-Hispanic<br>blacks | **0.03**<br>(0.01,0.06) | 0.02<br>(-0.01,0.05) | 0.02<br>(-0.01,0.05) | **0.04**<br>(0.01,0.07) | -0.04<br>(-0.09,0) |
| Non-Hispanic<br>Asian Americans | 0.03<br>(-0.01,0.08) | **0.04**<br>(0.01,0.06) | **0.12**<br>(0.08,0.17) | **0.06**<br>(0.01,0.11) | -0.03<br>(-0.09,0.03) |
| Non-Hispanic<br>Native Americans | **0.23**<br>(0.07,0.4) | **0.41**<br>(0.18,0.64) | **0.26**<br>(0.08,0.44) | **0.23**<br>(0.08,0.37) | -0.05<br>(-0.26,0.17) |

**Table 2:** Annual change in mortality rates involving poisoning by medicaments but not narcotics and psychodysleptics (ICD-10 codes T36-39.xx or T41-50.8 but not T40.xx not present as either underlying or contributing causes of death; category C in Table 1) between 2000-2011 and between 2011-2017 ...
| **2011-2017 period** |  |  |  |  |  |
| --- | --- | --- | --- | --- | --- |
|  | Ages<br>25-34y | Ages<br>35-44y | Ages<br>45-54y | Ages<br>55-64y | Ages<br>65-74y |
| Non-Hispanic<br>whites | **0.32**<br>**(0.22,0.42)** | **0.50**<br>**(0.41,0.59)** | **0.58**<br>**(0.45,0.7)** | **0.50**<br>**(0.41,0.58)** | **0.10**<br>**(0.05,0.16)** |
| Hispanics | **0.24**<br>**(0.19,0.29)** | **0.26**<br>**(0.19,0.33)** | **0.28**<br>**(0.17,0.4)** | **0.30**<br>**(0.22,0.38)** | 0.03<br>(-0.06,0.12) |
| Non-Hispanic<br>blacks | **0.25**<br>**(0.19,0.31)** | **0.47**<br>**(0.37,0.58)** | **0.38**<br>**(0.29,0.46)** | **0.28**<br>**(0.21,0.36)** | 0.02<br>(-0.03,0.06) |
| Non-Hispanic<br>Asian Americans | 0.03<br>(-0.01,0.06) | 0.08<br>(-0.04,0.2) | **0.19**<br>**(0.12,0.25)** | **0.27**<br>**(0.12,0.41)** | 0.02<br>(-0.08,0.12) |
| Non-Hispanic<br>Native Americans | **1.13**<br>**(0.87,1.39)** | **1.6**<br>**(1.12,2.08)** | **2.25**<br>**(1.67,2.83)** | **1.21**<br>**(0.64,1.78)** | 0.51<br>(-0.12,1.13) |

**Table 3:** Annual change in mortality rates involving poisoning by psychotropic drugs but not narcotics and psychodysleptics (ICD-10 codes T43.xx but not T40.xx present as either underlying or contributing causes of death; category D in Table 1) between 2000-2011 and between 2011-2017 per 100,000 individuals in different age/racial groups

| **2000-2011 period** |  |  |  |  |  |
| --- | --- | --- | --- | --- | --- |
|  | Ages<br>25-34y | Ages<br>35-44y | Ages<br>45-54y | Ages<br>55-64y | Ages<br>65-74y |
| Non-Hispanic<br>whites | **0.04**<br>**(0.01,0.06)** | **0.07**<br>**(0.03,0.1)** | **0.16**<br>**(0.13,0.19)** | **0.13**<br>**(0.11,0.15)** | **0.04**<br>**(0.03,0.05)** |
| Hispanics | **0.03**<br>**(0,0.05)** | 0.03<br>(0,0.07) | **0.08**<br>**(0.05,0.12)** | **0.07**<br>**(0.05,0.09)** | **0.04**<br>**(0.01,0.06)** |
| Non-Hispanic<br>blacks | 0.02<br>(-0.01,0.04) | 0.01<br>(-0.01,0.03) | **0.03**<br>**(0.01,0.05)** | **0.04**<br>**(0.02,0.06)** | -0.01<br>(-0.02,0.01) |
| Non-Hispanic<br>Asian Americans | 0.02<br>(0,0.05) | **0.04**<br>**(0.02,0.06)** | **0.11**<br>**(0.07,0.14)** | **0.06**<br>**(0.02,0.1)** | 0<br>(-0.04,0.05) |
| Non-Hispanic<br>Native Americans | **0.20**<br>**(0.05,0.34)** | **0.31**<br>**(0.13,0.5)** | **0.20**<br>**(0.08,0.33)** | **0.24**<br>**(0.08,0.4)** | 0.04<br>(-0.05,0.13) |

**Table 3:** Annual change in mortality rates involving poisoning by psychotropic drugs but not narcotics and psychodysleptics (ICD-10 codes T43.xx but not T40.xx present as either underlying or contributing causes of death; category D in Table 1) between 2000-2011 and between 2011-2017 per 100,000 ...
| **2011-2017 period** |  |  |  |  |  |
| --- | --- | --- | --- | --- | --- |
|  | Ages<br>25-34y | Ages<br>35-44y | Ages<br>45-54y | Ages<br>55-64y | Ages<br>65-74y |
| Non-Hispanic<br>whites | **0.31**<br>**(0.22,0.4)** | **0.52**<br>**(0.44,0.61)** | **0.59**<br>**(0.49,0.68)** | **0.49**<br>**(0.41,0.57)** | **0.14**<br>**(0.1,0.18)** |
| Hispanics | **0.22**<br>**(0.19,0.26)** | **0.29**<br>**(0.21,0.36)** | **0.24**<br>**(0.17,0.32)** | **0.29**<br>**(0.24,0.34)** | **0.10**<br>**(0.05,0.15)** |
| Non-Hispanic<br>blacks | **0.21**<br>**(0.18,0.25)** | **0.36**<br>**(0.27,0.45)** | **0.28**<br>**(0.21,0.35)** | **0.25**<br>**(0.19,0.32)** | **0.10**<br>**(0.05,0.15)** |
| Non-Hispanic<br>Asian Americans | 0.04<br>(0,0.09) | **0.11**<br>**(0.01,0.22)** | **0.20**<br>**(0.15,0.25)** | **0.26**<br>**(0.12,0.4)** | 0.04<br>(-0.06,0.14) |
| Non-Hispanic<br>Native Americans | **0.82**<br>**(0.4,1.25)** | **1.62**<br>**(1.3,1.95)** | **2.34**<br>**(1.81,2.87)** | **1.02**<br>**(0.62,1.42)** | **0.47**<br>**(0.05,0.89)** |

**Table 4:** Annual change in mortality rates involving poisoning by sedative-hypnotic drugs but not narcotics and psychodysleptics (ICD-10 codes T42.3-42.4 but not T40.xx present as either underlying or contributing causes of death; category E in Table 1) between 2000-2011 and between 2011-2017 per 100,000 individuals in different age/racial groups

| **2000-2011 period** |  |  |  |  |  |
| --- | --- | --- | --- | --- | --- |
|  | Ages<br>25-34y | Ages<br>35-44y | Ages<br>45-54y | Ages<br>55-64y | Ages<br>65-74y |
| Non-Hispanic<br>whites | **0.03**<br>**(0.02,0.04)** | **0.03**<br>**(0.02,0.04)** | **0.06**<br>**(0.05,0.06)** | **0.05**<br>**(0.04,0.06)** | 0.01<br>(0,0.02) |
| Hispanics | 0<br>(0,0.01) | **0.01**<br>**(0,0.02)** | **0.01**<br>**(0,0.02)** | 0.01<br>(0,0.03) | 0<br>(-0.01,0.01) |
| Non-Hispanic<br>blacks | **0.01**<br>**(0,0.01)** | 0.01<br>(0,0.01) | 0.01<br>(0,0.02) | 0<br>(-0.01,0.01) | 0<br>(-0.01,0.01) |
| Non-Hispanic<br>Asian Americans | 0.01<br>(-0.01,0.02) | **0.01**<br>**(0,0.02)** | 0.01<br>(0,0.03) | 0<br>(-0.03,0.02) | 0<br>(-0.02,0.02) |
| Non-Hispanic<br>Native Americans | 0.02<br>(-0.03,0.06) | **0.10**<br>**(0.04,0.16)** | **0.07**<br>**(0.04,0.09)** | -0.01<br>(-0.11,0.09) | -0.01<br>(-0.06,0.04) |

**Table 4:** Annual change in mortality rates involving poisoning by sedative-hypnotic drugs but not narcotics and psychodysleptics (ICD-10 codes T42.3-42.4 but not T40.xx present as either underlying or contributing causes of death; category E in Table 1) between 2000-2011 and between 2011-2017 ...
| **2011-2017 period** |  |  |  |  |  |
| --- | --- | --- | --- | --- | --- |
|  | Ages<br>25-34y | Ages<br>35-44y | Ages<br>45-54y | Ages<br>55-64y | Ages<br>65-74y |
| Non-Hispanic<br>whites | **0.06**<br>**(0.04,0.07)** | **0.05**<br>**(0.02,0.07)** | **0.06**<br>**(0.04,0.07)** | **0.05**<br>**(0.04,0.07)** | **0.03**<br>**(0.01,0.04)** |
| Hispanics | 0<br>(-0.01,0.01) | 0.01<br>(-0.01,0.03) | 0.01<br>(-0.03,0.05) | 0.02<br>(-0.03,0.06) | 0<br>(-0.02,0.01) |
| Non-Hispanic<br>blacks | **0.02**<br>**(0,0.04)** | **0.02**<br>**(0,0.04)** | **0.04**<br>**(0.02,0.07)** | 0.02<br>(-0.01,0.04) | 0<br>(-0.01,0.02) |
| Non-Hispanic<br>Asian Americans | 0.01<br>(-0.02,0.03) | -0.02<br>(-0.04,0.01) | 0.01<br>(-0.01,0.03) | 0.01<br>(-0.03,0.05) | 0.02<br>(-0.02,0.05) |
| Non-Hispanic<br>Native Americans | 0.21<br>(-0.03,0.45) | 0.05<br>(-0.17,0.28) | -0.07<br>(-0.24,0.09) | 0.04<br>(-0.05,0.13) | -0.05<br>(-0.24,0.15) |

Table 5 presents the estimates of annual increases in mortality rates involving poisoning by narcotics and psychodysleptics between 2000-2011, as well as between 2011-2017. Some of the largest increases in those rates are notably higher than the increases in the rates of mortality involving poisoning by medicaments but not narcotics and psychodysleptics (Table 5 vs. 2). For the 2000-2011 period, for younger age groups (25-54y), the largest increases in mortality rates involving poisoning by narcotics and psychodysleptics were in Non-Hispanic Native Americans and Non-Hispanic whites, with the corresponding rates of mortality in Hispanics and Non-Hispanic blacks decreasing for ages 35-44y; for older age groups (ages 55-74y), the largest increase between 2000-2011 was in Hon-Hispanic blacks. For the 2011-2017 period, for persons aged 25-44y, the largest increases in the rates of mortality involving poisoning by narcotics and psychodysleptics were in Non-Hispanic whites, followed by Non-Hispanic Native Americans/Non-Hispanic Blacks; for persons aged over 45y, the largest increases in the rates of mortality involving poisoning by narcotics and psychodysleptics between 2011-2017 were in Non-Hispanic blacks, followed by Non-Hispanic whites, then Non-Hispanic Native Americans. Increases in the rates of mortality involving poisoning by narcotics and psychodysleptics between 2011-2017 were lowest in Non-Hispanic Asians.

**Table 5:** Annual change in mortality rates involving poisoning by narcotics and psychodysleptics (ICD-10 codes T40.xx present as either underlying or contributing causes of death; category B in Table 1) between 2000-2011 and between 2011-2017 per 100,000 individuals in different age/racial groups

| **2000-2011 period** |  |  |  |  |  |
| --- | --- | --- | --- | --- | --- |
|  | Ages<br>25-34y | Ages<br>35-44y | Ages<br>45-54y | Ages<br>55-64y | Ages<br>65-74y |
| Non-Hispanic<br>whites | **1.24**<br>**(1.15,1.34)** | **0.76**<br>**(0.59,0.92)** | **1.17**<br>**(0.98,1.37)** | **0.78**<br>**(0.74,0.83)** | **0.19**<br>**(0.17,0.21)** |
| Hispanics | 0.07<br>(0,0.15) | **-0.28**<br>**(-0.38,-0.18)** | 0.01<br>(-0.19,0.21) | **0.33**<br>**(0.27,0.4)** | **0.08**<br>**(0.03,0.13)** |
| Non-Hispanic<br>blacks | -0.05<br>(-0.18,0.08) | **-0.73**<br>**(-1.08,-0.37)** | -0.1<br>(-0.64,0.43) | **1.01**<br>**(0.8,1.22)** | **0.23**<br>**(0.09,0.37)** |
| Non-Hispanic<br>Asian Americans | **0.10**<br>**(0.06,0.15)** | **0.04**<br>**(0,0.08)** | **0.05**<br>**(0,0.1)** | **0.08**<br>**(0.05,0.11)** | 0.01<br>(-0.02,0.04) |
| Non-Hispanic<br>Native Americans | **1.11**<br>**(0.84,1.39)** | **1.4**<br>**(0.85,1.96)** | **1.52**<br>**(1.18,1.85)** | **0.94**<br>**(0.69,1.2)** | 0.21<br>(-0.13,0.54) |
| **2011-2017 period** |  |  |  |  |  |
|  | Ages<br>25-34y | Ages<br>35-44y | Ages<br>45-54y | Ages<br>55-64y | Ages<br>65-74y |
| Non-Hispanic<br>whites | **4.34**<br>**(3.04,5.65)** | **3.74**<br>**(2.62,4.86)** | **2.1**<br>**(1.47,2.73)** | **1.79**<br>**(1.54,2.04)** | **0.44**<br>**(0.4,0.49)** |
| Hispanics | **1.35**<br>**(0.88,1.82)** | **1.24**<br>**(0.82,1.66)** | **0.86**<br>**(0.39,1.34)** | **0.81**<br>**(0.54,1.08)** | **0.40**<br>**(0.28,0.52)** |
| Non-Hispanic<br>blacks | **2.05**<br>**(1.22,2.87)** | **2.64**<br>**(1.61,3.67)** | **3.39**<br>**(1.76,5.03)** | **4.00**<br>**(2.53,5.47)** | **2.3**<br>**(1.74,2.86)** |
| Non-Hispanic<br>Asian Americans | **0.46**<br>**(0.31,0.6)** | **0.21**<br>**(0.05,0.37)** | **0.15**<br>**(0.03,0.27)** | 0.08<br>(-0.01,0.16) | 0.04<br>(-0.04,0.12) |
| Non-Hispanic<br>Native Americans | **2.22**<br>**(1.2,3.24)** | **2.14**<br>**(0.54,3.74)** | **1.15**<br>**(0.25,2.04)** | **0.93**<br>**(0.38,1.47)** | 0.34<br>(-0.19,0.88) |

Table 6 presents the relative (fold) increases in the average annual rates of mortality for categories B-E in Table 1 in different age groups in the overall population of US adults for the 2015-2017 period vs. 2000-2002 period, as well as for the 2011-2014 period vs. 2000-2002 period. Table 6 shows major increases in mortality involving poisoning, particularly by narcotics and psychodysleptics (especially in persons aged 55-74y), as well as psychotropic medications, particularly for individuals aged 45-64y. Increases in the rates of mortality involving poisoning by both psychotropic medications but not narcotics and psychodysleptics, as well as for sedative-hypnotic medications but not narcotics and psychodysleptics for the 2011-2014 period vs. 2000-2002 period were greater than the corresponding increases in the rates of prescribing for those medications, particularly for persons aged 45-64y (Table 6 vs. [31]) – see Discussion.

**Table 6:** Relative (fold) increases in the rates of mortality involving poisoning by different substances (categories C, D, E, and B in Table 1) for different age groups of US adults for the 2015-2017 period vs. the 2000-2002 period, as well as the 2011-2014 period vs. the 2000-2002 period.

| 2015-2017 vs. 2000-2002 periods | Ages 25-34y | Ages 35-44y | Ages 45-54y | Ages 55-64y | Ages 65-74y |
| --- | --- | --- | --- | --- | --- |
| medicaments but not narcotics and psychodysleptics | 2.33 | 2.07 | 2.56 | 3.1 | 1.43 |
| psychotropic drugs but not narcotics and psychodysleptics | 3.22 | 2.75 | 3.48 | 5.37 | 3.94 |
| sedative-hypnotic drugs but not narcotics and psychodysleptics | 3.06 | 1.75 | 2.27 | 2.75 | 1.73 |
| narcotics and psychodysleptics | 4.29 | 2.32 | 2.62 | 6.54 | 6.33 |
| 2011-2014 vs. 2000-2002 periods | Ages 25-34y | Ages 35-44y | Ages 45-54y | Ages 55-64y | Ages 65-74y |
| medicaments but not narcotics and psychodysleptics | 1.64 | 1.48 | 1.92 | 2.24 | 1.27 |
| psychotropic drugs but not narcotics and psychodysleptics | 1.95 | 1.65 | 2.3 | 3.24 | 2.5 |
| sedative-hypnotic drugs but not narcotics and psychodysleptics | 2.26 | 1.47 | 2 | 2.31 | 1.35 |
| narcotics and psychodysleptics | 2.42 | 1.38 | 1.89 | 4.14 | 3.82 |

There were no correlations between state-specific rates of mortality involving poisoning by psychotropic drugs but not narcotics/psychodysleptics and state-specific rates of mortality involving poisoning by narcotics/psychodysleptics during the 2013-2017 period, suggesting potential differences in factors behind those two epidemics; those correlations were −0.06 (95% CI (−0.33,0.22)) for persons aged 45-64y, and −0.16(−0.42,0.12) for persons aged 25-44y. Further details on the geographic spread of the two epidemics are presented in the Supporting Information.

## Discussion

While the rise in mortality by narcotic, including opiate poisoning, particularly during the most recent years in well documented ([5,9]), our understanding of the contribution of medicaments other than narcotics to mortality is more limited. Moreover, there are differences in the rates of consumption of different medications between the different racial groups [10–18], and those differences may reflect differences in the rates of adverse outcomes associated with medication use in different racial groups [19–24;26,27]. In this study, we have shown significant increases in the rates of mortality involving poisoning by medicaments other than narcotics and psychodysleptics in different age/racial groups, both for the 2000-2011 period, and even more so for the 2011-2017 period. Those increases were particularly large in non-elderly Non-Hispanic Native Americans and whites. The majority of deaths involving poisoning by medicaments other than narcotics and psychodysleptics, as well as most of the increases in the corresponding mortality rates were for deaths involving poisoning by psychotropic medications.

Significant increases in prescribing for antidepressants and antipsychotic (as well as sedative-hypnotic) drugs in different age groups of US adults took place during the study period [31]. For example, for the 2011-2014 period, 17.5% of US adults aged 45-64y and 18.7% of those aged 65-74y reported receiving antidepressant medications during the last 30 days, compared to 10.5% and 9.3% correspondingly for the 1999-2002 period. At the same time, relative increases in the rates of mortality involving poisoning by psychotropic medications but not narcotics and psychodysleptics were greater than increases in prescribing for psychotropic medications, particularly for persons aged 45-64y (Table 6 vs. [31]). This means that the likelihood of a poisoning death per unit of prescribing of psychotropic or sedative-hypnotic medications has increased in the overall population of US adults to whom those medications are prescribed, particularly in the 45-64y age group. This doesn’t appear to be the case in several other countries (e.g. Australia), where the rates of death per unit of prescribing of antipsychotic medication have decreased somewhat with time [32], which may be related to the shift to less toxic drugs [33,32]. In the UK and the European union, rates of mortality involving poisoning by drugs and medications are notably lower than in the US [34,35]; moreover, in the UK, rates of mortality involving poisoning by both narcotics and antipsychotic drugs have not experienced such major increases as the corresponding rates during the same period in the US [35]. Further work is needed to better understand the types of individuals, as well as the types of drugs/medications (e.g. 1^st^ vs. 2^nd^ generation antipsychotic medications [33,32]) that are associated with fatal outcomes resulting from consumption of those substances. We should note that while rates of prescribing of certain medications such as antidepressants to US adults aged over 75y have increased significantly [31], rates of mortality involving poisoning by medications but not narcotics/psychodysleptics in US adults aged over 75y have decreased with time (results not included), possibly having to do with a shift to less toxic drugs [33]. Comparison of the mortality trends between US adults aged over 75y vs. non-elderly US adults points to the possibility of self-harm being a factor in the increase in the rates of mortality associated with poisoning by medications in non elderly US adults (particularly non-Hispanic whites), which may also be related to the increases in the rates of mortality associated with alcohol poisoning and suicide in those population groups [8].

While rates of mortality involving poisoning by medicaments other than narcotics and psychodysleptics are lower than the rates of mortality involving poisoning by narcotics and psychodysleptics, the pernicious effects that medication use/misuse may have on health, including mortality outcomes are not restricted to deaths that involve poisoning by medicaments listed on a death certificate. Indeed, a number of studies, e.g. [19–24] have suggested elevated risks for mortality and other health-related outcomes associated with the use of psychotropic medications (both antipsychotic medications and antidepressants) and sedative-hypnotic medications (particularly benzodiazepines). Recent work suggests associations between rates of prescribing for penicillins, and rates of mortality and hospitalization for septicemia/sepsis in older adults [26,27]. Additionally, rates of consumption of certain medications, particularly psychotropic drugs, sedative-hypnotic medications and antibiotics in Non-Hispanic whites are higher than in other major racial groups (save for Native Americans) [10–18], and those differences may have an effect on health-related, including mortality outcomes [19–24,26,27]. Our results, as well as other work, e.g. [19–24,26,27] support the need for a comprehensive evaluation of the impact of prescribing/misuse of various medications on health, including mortality outcomes.

Our paper has some limitations. It might be that increases in the rates of mortality involving poisoning by medicaments other than narcotics were affected by changes in notification and coding practices. For example, in light of the increases in the rates of mortality involving poisoning by narcotics/psychodysleptics, a growing fraction of deaths having poisoning by medicaments but not poisoning by narcotics/psychodysleptics on a death certificate could have actually been partly affected by narcotic/psychodysleptic use, with poisoning by narcotics/psychodysleptics not appearing on some of those death certificates. We do not believe that this could largely explain our results as there are no correlations between state-specific rates of mortality involving poisoning by psychotropic drugs but not narcotics/psychodysleptics and state-specific rates of mortality involving poisoning by narcotics/psychodysleptics during the 2013-2017 period (Results; Supporting Information). Additionally, there are notable discrepancies in trends for mortality involving poisoning by narcotics and psychodysleptics vs. mortality involving poisoning by medicaments other than narcotics, particularly for Hispanics and Non-Hispanic blacks, further supporting the notion that deaths involving poisoning by narcotics and psychodysleptics and deaths involving poisoning by medicaments other than narcotics represent different epidemics, and increases in the recorded rates of the latter deaths are not an artifact of the increases in the rates of the former deaths. For example, rates of mortality involving poisoning by medicaments but not narcotics/psychodysleptics in Hispanics aged 35-44y increased between 2000-2011 (Table 2 and Figure 3), while rates of mortality involving poisoning by narcotics and psychodysleptics in that population declined between 2000-2011 (Table 5 and Figure 2); for persons aged 55-64y, increases in mortality involving poisoning by narcotics and psychodysleptics were greater in Non-Hispanic blacks than in Non-Hispanic whites for both the 2000-2011 and the 2011-2017 periods (Table 5), with the opposite being true for increases in mortality involving poisoning by medicaments but not narcotics and psychodysleptics (Table 2). Another possible contributing factor to the increase in the rates of mortality involving poisoning by medicaments but not narcotics and psychodysleptics is change in diagnostic/coding criteria. However, it is unlikely that such changes, if they indeed took place, would be very different for different groups of adults, while growth in the rates of mortality involving poisoning by medicaments but not narcotics and psychodysleptics is notably higher for some age/racial groups compared to others. For example, there were no discernible changes in the rates of mortality involving poisoning by medicaments but not narcotics and psychodysleptics in Non-Hispanic blacks aged 34-54y between 2000-2011 (Table 2), while major changes in the corresponding mortality rates took place in Non-Hispanic whites and Non-Hispanic Native Americans. Another example is Non-Hispanic whites aged 55-64y vs. Non-Hispanic whites aged 65-74y. In the beginning of the study period, rates of mortality involving poisoning by medicaments but not narcotics and psychodysleptics were similar for the two age groups; however, subsequently those rates followed quite different trends for the two age groups (Figure 3). We believe that those differences in trends for different population groups are unlikely to be explained by differences in changes in diagnostic/coding criteria for those population groups, but rather by genuine differences in mortality trends between the different population groups. Finally, we don’t know whether increases in prescribing or misuse have contributed to the increases in the rates of mortality involving poisoning by medicaments but not narcotics and psychodysleptics. Significant increases in medication prescribing rates, particularly for antipsychotic and sedative-hypnotic drugs in persons aged 25-44y, and antidepressants in persons aged 45-74y did take place during the study period [31]. At the same time, increases in mortality involving poisoning by psychotropic, as well as sedative-hypnotic drugs, but not narcotics and psychodysleptics were generally greater than increases in prescribing for those medications, particularly for persons aged 45-64y (compare Table 6 to [31]). Additionally, while rates of prescribing of psychotropic medications in individuals aged over 75y increased during the study period [31], rates of mortality involving poisoning by those medications in individuals aged over 75y have decreased, which is unlike the case of younger individuals. All of this suggests the potential role of misuse of the antidepressant/antipsychotic, as well as sedative-hypnotic medications in the increases in the rates of mortality by poisoning in certain population groups, particularly non-elderly white and Native Americans.

We believe that despite some limitations, our study shows significant increases in the rates of mortality involving poisoning by medicaments but not narcotics and psychodysleptics, especially poisoning by psychotropic drugs in different age/racial groups, particularly non-elderly Non-Hispanic Native Americans and whites, both between 2000-2011, and even more so between 2011-2017. Moreover, some of those increases in mortality rates have significantly exceeded the increases in prescription rates for the corresponding medications, pointing to the potential role of medication misuse. Those results, as well as earlier studies showing associations between medication use and health-related and mortality outcomes (e.g. [19–24,26,27]), including for deaths that do not have poisoning listed on a death certificate further support the need for a comprehensive evaluation of the impact of prescribing and misuse of various medications on health and mortality outcomes in different population groups, as well as the related guidance regarding the use of those medications. Additionally, such analyses may help elucidate the role of medication use/misuse in the long-term decline in life expectancy in Non-Hispanic whites compared to other major racial groups in the US [1].

---

Supporting Information for “Rise in mortality involving poisoning by medicaments other than narcotics, including poisoning by psychotropic drugs in different age/racial groups in the US”.

**Section S1**: Spatial variability in the rates of mortality involving poisoning by psychotropic medications other narcotics and psychodysleptics, and rates of mortality involving poisoning by narcotics and psychodysleptics

As suggested in the Results section in the main body of the text, there were no correlations between state-specific rates of mortality involving poisoning by psychotropic drugs but not narcotics/psychodysleptics (category D in Table 1) and state-specific rates of mortality involving poisoning by narcotics/psychodysleptics (category B in Table 1) during the 2013-2017 period; those correlation were −0.06(95% CI (−0.33,0.22)) for persons aged 45-64y, and −0.16(−0.42,0.12) for persons aged 25-44y. Table S1 lists the 20 states with the highest rates of mortality involving poisoning by psychotropic drugs but not narcotics/psychodysleptics, as well as the 20 states with the highest rates of mortality involving poisoning by narcotics/psychodysleptics per 100,000 persons aged 25-44y during the 2013-2017 period; Table S2 presents the corresponding lists for persons aged 45-64y, with all the data extracted from [S1]. Lists of states with the highest rates of mortality involving poisoning by psychotropic drugs but not narcotics/psychodysleptics are quite different from lists of states with the highest rates of mortality involving poisoning by narcotics/psychodysleptics, with a notable presence of western states among those with the highest rates of mortality involving poisoning by psychotropic drugs but not narcotics/psychodysleptics (particularly for persons aged 45-64y) vs. a notable presence of eastern (particularly northeastern) states among those with the highest rates of mortality involving poisoning by narcotics/psychodysleptics. Furthermore, rates of mortality involving poisoning by psychotropic drugs but not narcotics/psychodysleptics are higher for persons aged 45-64y compared to persons aged 25-44y, while rates of mortality involving poisoning by narcotics/psychodysleptics are somewhat higher for persons aged 25-44y compared to persons aged 45-64y.

**Table S1:**
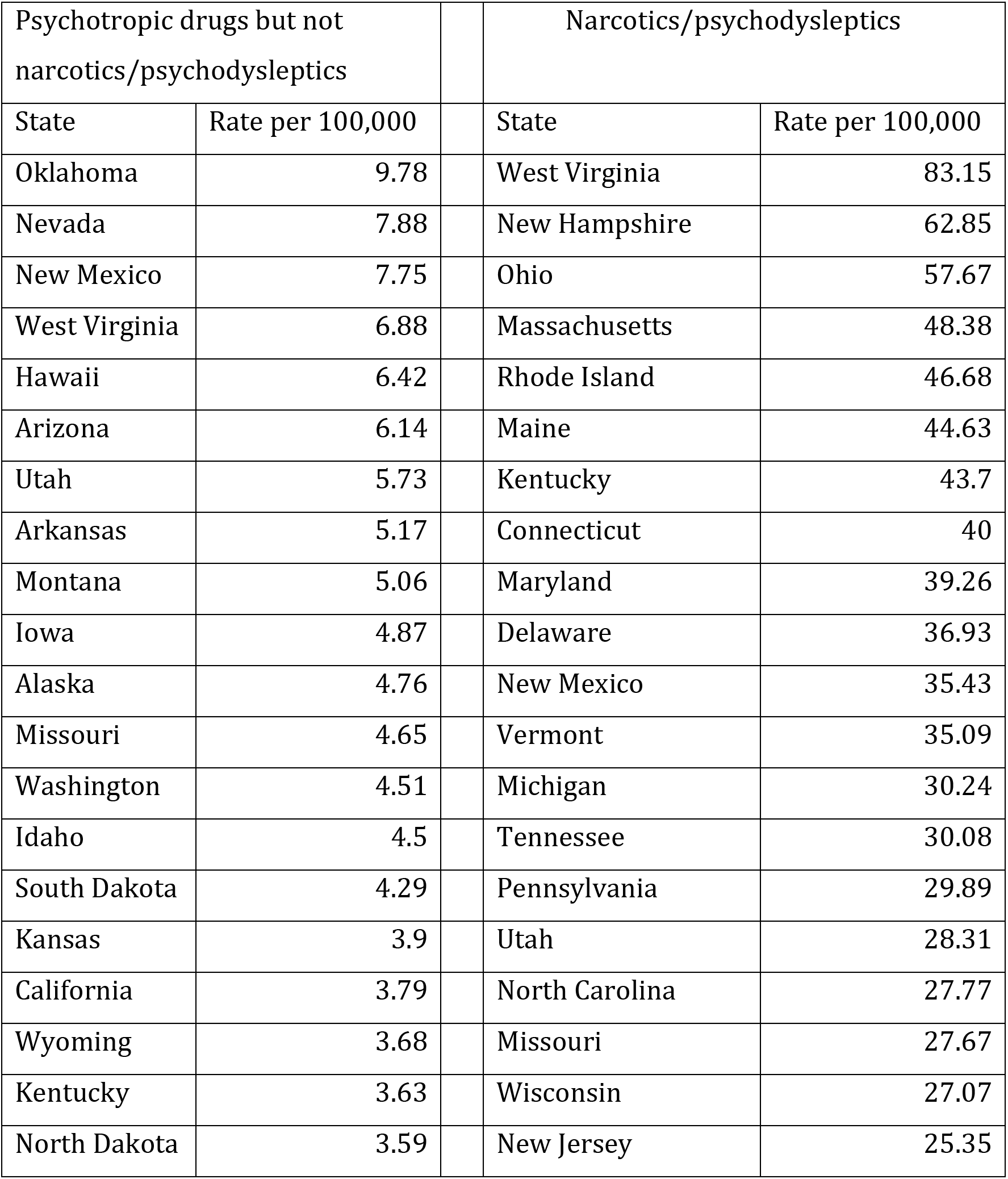
20 states with the highest rates of mortality involving poisoning by psychotropic drugs but not narcotics/psychodysleptics, as well as the 20 states with the highest rates of mortality involving poisoning by narcotics/psychodysleptics per 100,000 persons aged 25-44y during the 2013-2017 period (including the corresponding mortality rates).

| Psychotropic drugs but not narcotics/psychodysleptics |  | Narcotics/psychodysleptics |  |
| --- | --- | --- | --- |
| State | Rate per 100,000 | State | Rate per 100,000 |
| Oklahoma | 9.78 | West Virginia | 83.15 |
| Nevada | 7.88 | New Hampshire | 62.85 |
| New Mexico | 7.75 | Ohio | 57.67 |
| West Virginia | 6.88 | Massachusetts | 48.38 |
| Hawaii | 6.42 | Rhode Island | 46.68 |
| Arizona | 6.14 | Maine | 44.63 |
| Utah | 5.73 | Kentucky | 43.7 |
| Arkansas | 5.17 | Connecticut | 40 |
| Montana | 5.06 | Maryland | 39.26 |
| Iowa | 4.87 | Delaware | 36.93 |
| Alaska | 4.76 | New Mexico | 35.43 |
| Missouri | 4.65 | Vermont | 35.09 |
| Washington | 4.51 | Michigan | 30.24 |
| Idaho | 4.5 | Tennessee | 30.08 |
| South Dakota | 4.29 | Pennsylvania | 29.89 |
| Kansas | 3.9 | Utah | 28.31 |
| California | 3.79 | North Carolina | 27.77 |
| Wyoming | 3.68 | Missouri | 27.67 |
| Kentucky | 3.63 | Wisconsin | 27.07 |
| North Dakota | 3.59 | New Jersey | 25.35 |

**Table S2:**
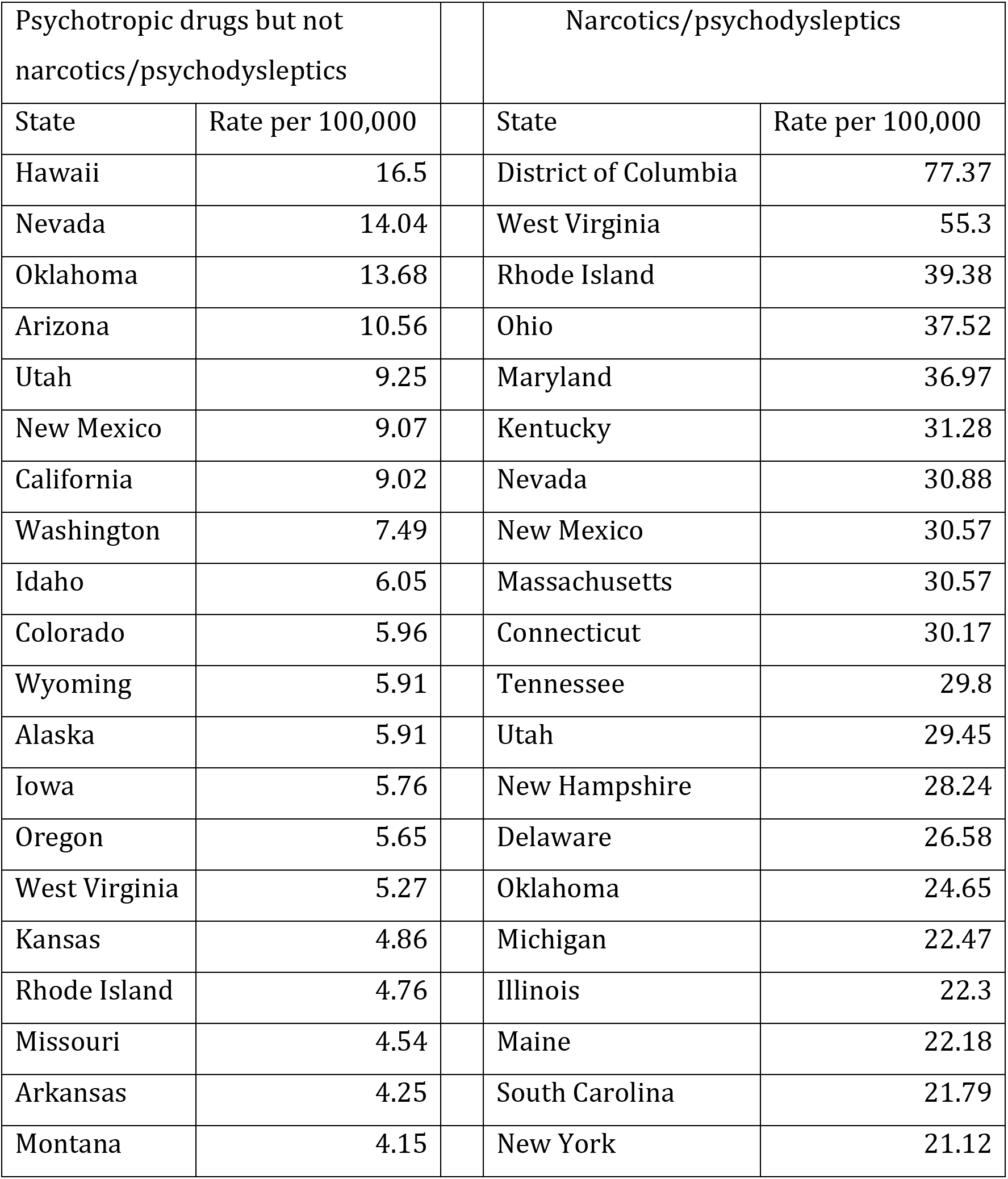
20 states with the highest rates of mortality involving poisoning by psychotropic drugs but not narcotics/psychodysleptics, as well as the 20 states with the highest rates of mortality involving poisoning by narcotics/psychodysleptics per 100,000 persons aged 45-64y during the 2013-2017 period (including the corresponding mortality rates).

| Psychotropic drugs but not narcotics/psychodysleptics |  | Narcotics/psychodysleptics |  |
| --- | --- | --- | --- |
| State | Rate per 100,000 | State | Rate per 100,000 |
| Hawaii | 16.5 | District of Columbia | 77.37 |
| Nevada | 14.04 | West Virginia | 55.3 |
| Oklahoma | 13.68 | Rhode Island | 39.38 |
| Arizona | 10.56 | Ohio | 37.52 |
| Utah | 9.25 | Maryland | 36.97 |
| New Mexico | 9.07 | Kentucky | 31.28 |
| California | 9.02 | Nevada | 30.88 |
| Washington | 7.49 | New Mexico | 30.57 |
| Idaho | 6.05 | Massachusetts | 30.57 |
| Colorado | 5.96 | Connecticut | 30.17 |
| Wyoming | 5.91 | Tennessee | 29.8 |
| Alaska | 5.91 | Utah | 29.45 |
| Iowa | 5.76 | New Hampshire | 28.24 |
| Oregon | 5.65 | Delaware | 26.58 |
| West Virginia | 5.27 | Oklahoma | 24.65 |
| Kansas | 4.86 | Michigan | 22.47 |
| Rhode Island | 4.76 | Illinois | 22.3 |
| Missouri | 4.54 | Maine | 22.18 |
| Arkansas | 4.25 | South Carolina | 21.79 |
| Montana | 4.15 | New York | 21.12 |

